# LSD1 demethylase inhibition prevents cardiac fibrosis in both ischemic and congenital diseases in mice and pig models

**DOI:** 10.1101/2024.12.08.627384

**Authors:** Denis Arnaud, Virginie Lambert, David Benoîst, Amandine Martin, Jean Gauthier, Michel Pucéat

## Abstract

Fibrosis is part of a clinical burden in cardiovascular diseases. The pathological process has been the subject of intensive research with still mitigated therapeutic options. Recently chromatin modifiers have turned out to be potential drugs to modulate fibrosis. Here, in order to address the question of pharmacological inhibition of fibrosis, we used both a mouse model of myocardial infarction with left ventricular fibrosis and a more clinically relevant pig model of right ventricular failure featuring interstitial fibrosis. Treatment of these diseased animal models with an inhibitor of the Lysine Demethylase 1 (LSD1) significantly prevented both left and right ventricular failure in both the mouse and the pig, respectively. This was revealed by a significant recovery of left ventricular function post-myocardial infarction in the mouse and a limitation of remodeling of the pig right ventricle, thus preserving its function. Fibrosis was significantly decreased in both mouse and pig hearts, which likely account for improvement in ventricular function. We thus provide evidence of the beneficial effect of LSD1 inhibitors in cardiac fibrosis and of the use of such drugs to preserve ventricular function in both ischemic and congenital heart diseases.

**NEW & NOTEWORTHY:** Drugs to prevent cardiac fibrosis has been the subject of intensive research with limited outcomes. This work inspired by oncology, provides evidence that an epigenetic modifier which targets the process of epithelial-to-mesenchymal transition turns out to be an efficient inhibitor of fibrosis for both ischemic and non-ischemic myocardial diseases.

## INTRODUCTION

Cardiovascular diseases remain responsible for a large majority of morbidity and mortality in the vast majority of the world (1). In both ischemic diseases mostly affecting the left ventricle and in congenital diseases with right ventricular failure often due to ventricular overload and/or overpressure, myocytes cannot ensure any longer their function and die. The death of diseased myocytes is compensated by fibroblasts which when activated together with pericytes rapidly convert into myofibroblasts. The later express and secrete many proteins of the extracellular matrix including smooth muscle actin, periostin, PDGF receptors and, collagen 1a1, and fibronectin, which triggers a pathophysiological process so called fibrosis(2). First adaptative, fibrosis then leads to severe functional defects including deficient contractility and conduction disorders.

Fibrosis often represents the final stage of numerous diseases including cardiovascular, lung and kidney diseases. Despite the tremendous literature on fibrosis, few therapies have been approved at present to target cardiac fibrosis (3).

Recently, studies including epigenetics approaches, uncovered bromo domain and extra-terminal domain (BET) inhibitors as potential anti-fibrotic agents(4, 5) through a TGF dependent manner(6), a pathway targeted by inhibitors or antibodies in many clinical trials with still mitigated results.

Acetylation of proteins has been also proposed to play a role in cardiac fibrosis(7) making deacetylase potential drugs to alleviate fibrosis. More recently epitranscritomics has been involved in the fibrosis process and related to epigenetics(8). A very recent paper reported that an HDAC inibitor used in a mouse model of myocardial infarction alleviated inflammation and fibrosis through an inhibition of LSD1(9)

A novel approach largely used in oncology is emerging for cardiac fibrosis: the CAR T-cell therapy. Fibroblast Activated Protein (FAP) being among the most upregulated membrane protein in activated fibroblast, FAP CAR T-cells were engineered and infused to mouse injured heart. They could decrease fibrosis(10, 11).

Interestingly, biological pathways targeted in the treatment of cardiac fibrosis are also targeted in oncology to limit metastasis. Both pathological phenomenon share the same biological process: namely the epithelio-to-mesenchymal transition (EMT).

The epigenetic modifier LSD1 (lysine-specific demethylase 1) plays a crucial role in the formation of metastasis and LSD1 inhibitor prevents the phenomenon by blocking EMT(12). This leads to the development of many LSD1 inhibitors used in clinical trials to target metastasis(13).

TGFβ-induced fibrosis in skin, lung, liver, and kidney in mouse models was blocked by the formation of LSD1/NR4A1 complex on profibrotic gene regulatory modules(14).

We also reported that inhibition of LSD1 at neonatal stage, prevented fibrosis in a genetic mouse model of dilated cardiomyopathy, i.e., a laminopathy(15).

LSD1 expression is increased in patients featuring dilated cardiomyopathy as well as in mouse model of pressure overload-induced fibrosis. Knock-down of LSD1 in fibroblasts prevented fibrosis and improved ventricular function in a mouse model(16).

Here, we tested LSD1 inhibitors on the development of fibrosis in both mouse models of left myocardial infarction and in a more clinically relevant pig model of right ventricular failure. Using tissue histology and echocardiography, we found that LSD1 inhibitors significantly prevented fibrosis and improved ventricular function in both mice and pigs.

## MATERIALS AND METHODS

### Mice

6-8 weeks mice (OF1) were subjected to surgery to perform left coronary ligation.

Mice were anesthetized using a mixture of ketamine (71.5 mg/kg) and xylazine (14.5 mg/kg) via intraperitoneal injection. Bupremorphine (0.1 mg/Kg) and carprofen (5 mg/Kg) were given as analgesic. The chest wall was shaved. Drops of lidocaine (21.33 mg/ml) were added on the skin before opening the chest.

Surgery to lead to permanent ( occlusion of the artery with a definite ligature) or transient ( ligature released after 30 min) left coronary artery was performed as previously published(17).

Atipamezole (0.5mg/Kg), sulfadomethoxin (30 mg/Kg) were given post-surgery.

The mice were left for recovery for 10 days post-surgery and then treated with GSK-LSD1 (1.5 mg/kg), a reversible LSD1 inhibitor or losartan (5 mg/Kg) by gavage for 3 weeks 3 times/week. SP- 2509 a reversible LSD1 inhibitor was also used at 30 mg/Kg.

Mice were weighted every day. No difference of weight was observed between groups (Supplementary Figure 1).

### Doppler ultrasound transthoracic echocardiography

Echocardiography was performed 5 weeks post-surgery using an Affiniti 50 (Philips) and a 18-MHz probe. Mice were anesthetized with 4% isoflurane and maintained with 1 to 1.5% isoflurane. Ventricular contractility was monitored using M-mode by positioning the M-mode cursor within the mitral inflow stream pre-recorded by Doppler mode.

### Histology and immunofluorescence analysis

After heart explantation from pigs, several around 1 cm3 blocks were cut from the free wall of the right ventricle and were freeze-clamped in liquid nitrogen. Blocks embedded in OCT were then cut in 10 μm sections. Different sections from each block were fixed in PFA 4% and then stained. Adult mouse were fixed in 4% paraformaldehyde overnight at 4°C, then dehydrated with a series of ethanol washes, and embedded in paraffin. The blocks were sectioned at a thickness of 8 μm. The tissue was rehydratated and sections were stained with eosin and hematoxylin and Sirius Red (collagen) according to the manufacturer’s instructions. At least 100 images were acquired with a scanner (Zeiss axioscan). Staining intensity and area of red staining were measured using ImageJ (NIH, Bethesda, MD) for histology images and IMARIS (Oxford instruments) for immunosfluorescence images. We first used the ImageJ plugin Color Deconvolution to separate channels. Then the images were converted into grays scale images and inverted. The stained areas were then segmented using thresholding. Thresholded pixels were normalized to foreground color. The segmented areas were then measured.

Mouse heart sections were also treated with xylene and alcohol to remove paraffin, and after a step of antigen retrieval were stained for anti-smooth muscle actin (SMA), anti-troponin T (Tnt) and dapi (DNA). SMA+ cells were scored using IMARIS software in at least 3 sections per heart. In IMARIS, we first selected the channel used for SMA staining. Then we used the surface tool and next the mask option (thresholding using the image display) which created a new masking channel. The surface of interest was then displayed in the statistical tab.

### Pig surgery

All experiments were carried out according to the European Community guiding principles in animal experimentation. Four piglets underwent surgery mimicking repaired tetralogy of Fallot as previously described(18). Under general anesthesia, through a left thoracotomy approach, a side-biting vascular clamp was longitudinally set across the pulmonary valve annulus without obstruction of the RV outflow tract. A pulmonary valve leaflet was excised and the pulmonary infundibulum, annulus, and trunk were enlarged by a 2-cm-long, elliptically shaped polytetrafluoroethylene patch to ensure the loss of valve integrity. This chronic pulmonary valve regurgitation led to RV volume overload. Branch pulmonary arterial obstruction is not uncommon after surgical correction of TOF related to native hypoplasia or surgical complications and is known to increase pulmonary valve regurgitation. In our model, RV pressure overload was achieved by pulmonary artery banding, made of umbilical tape, placed around the artery truncus and secured for a final diameter of approximately 1 cm to ensure progressive pulmonary stenosis with animal maturation. After 5 months (median: 158 ± 48 days), the combined pressure and volume overloaded RV leads to its contractile failure. Pigs were treated with the LSD1 inhibitor (1.5 mg/Kg) incorporated in the food 3 weeks (median 21 days) after the surgery for a median duration of 30 days.

### Pig heart echocardiography

RV function was evaluated in all animals at each step of experimental procedure at baseline before rTOF surgery, at and at 5 months at the end of study.

Echocardiography was performed on closed-chest animals under general anesthesia in the dorsal decubitus position. We used a commercially available Vivid E9 ultrasound machine (GE Medical Systems, Milwaukee, WI) equipped with a 2.5-MHz transducer. The values of all echocardiographic parameters were obtained as the average value of three consecutive cardiac cycles during transient apnea and analyzed on a comprehensive workstation (EchoPAC 110.1.2; GE Vingmed Ultrasound, Horten, Norway). Standard parameters of RV function such as Fraction of Area Change, Tricuspid Annular Systolic Plane Excursion (TAPSE), S’wave on TDI) as well as myocardial strain (longitudinal global and free wall strain) were monitored.

#### LSD1 enzymatic activity

LSD1 was assayed using a kit (Epigenetics) according to the manufacturer.

#### Statistics

Data were analyzed by Graphpad Prism using two-ways anova or T-tests.

## RESULTS

We first used a mouse model of myocardial infarction (MI) following permanent or transient left coronary ligature. Six to eight months old mice were treated with 1.5mg GSK-LSD1/Kg or with 5mg/Kg Losartan 10 days post-surgery for 3 weeks, 3 days/week. Echocardiography was performed before surgery, ten days post-surgery and after 3 weeks to assess left ventricular function. Mice were then euthanized. The heart was explanted, and sectioned for histological monitoring.

Figure 1A shows that myocardial infarction was severe. The ventricular wall was significantly thinner in mice subjected to MI than under negative control condition (defined as healthy mice with no MI and no treatment) as revealed by eosin/hematoxylin staining. Sirius red staining also revealed an extended fibrosis in mice with MI while it was absent in control mice. While Losartan partially limited the extent of fibrosis (Figure 1A,B) in mice subjected to MI, GSK-LSD1 that diminished the enzymatic activity of the demethylase (Supplementary Figure 2) even significantly further decreased the impact of the pathophysiological process. This was accompanied by a thickening of the left ventricular wall that was more prominent in GSK-LSD1- treated mice than in losartan-treated mice (Figure 1C).The reversible LSD1 inhibitor SP 2509 used at 30mg/Kg also significantly decreased fibrosis in the post-MI hearts to the same extent as GSK-LSD1 (Supplementary Figure 3).

**Figure 1.**
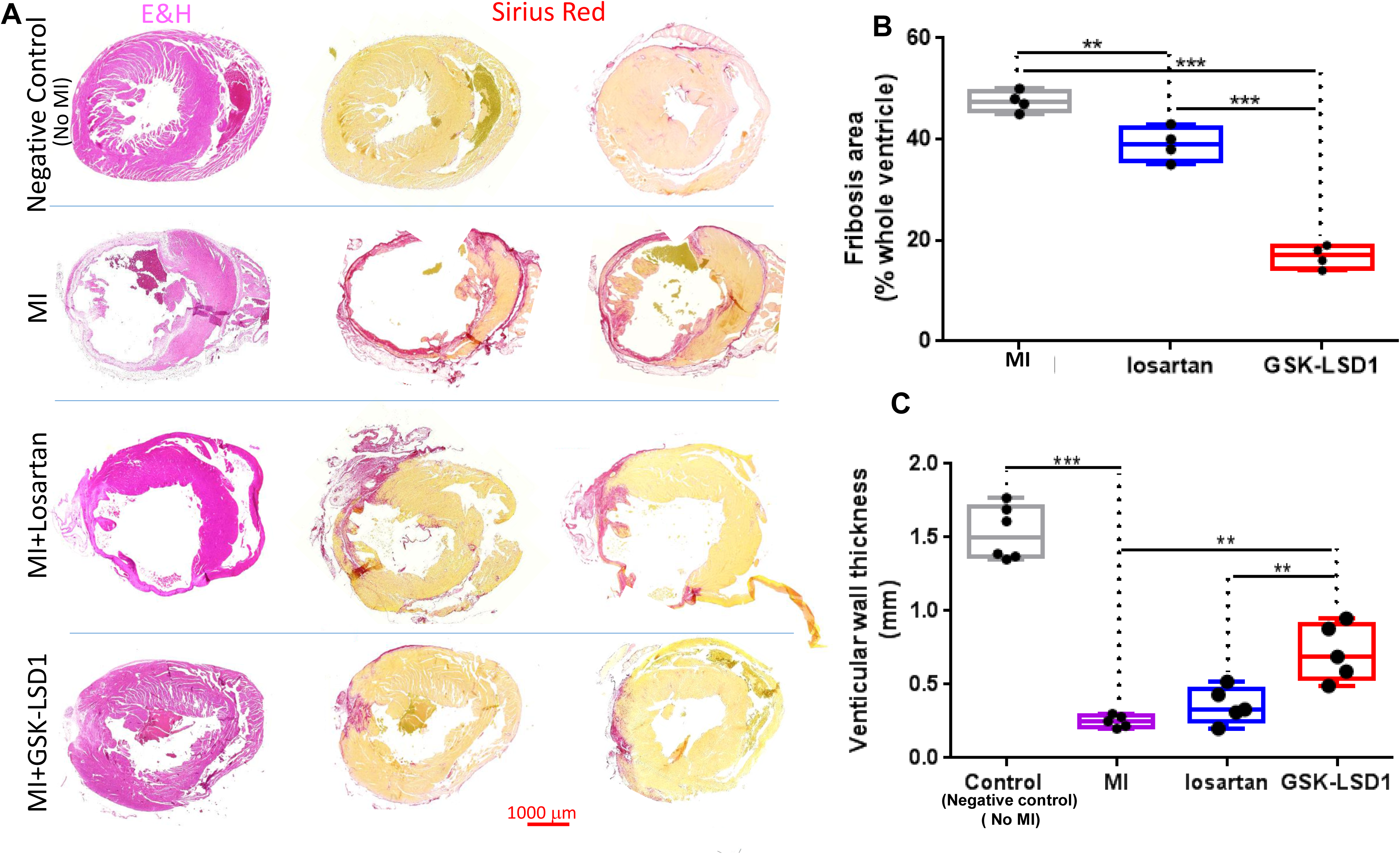
Fibrosis in mouse hearts. **(A)** eosin/hematoxylin (E&H) (left panels) and sirius red (right 2 panels) of heart sections under different experimental conditions, negative controls were healthy mice with no MI and no treatment (B) extent of fibrosis and (C) left ventricular wall thickness in MI, losartan- and GSK-LSD1-treated mouse hearts. Significantly different ** p< 0,01,***p<0,001, n=5 mice.

Ventricular function was assessed by echocardiography. Figure 2 shows that the MI-induced decrease in shortening fraction was sligthly but not significantly improved by Losartan while contractility was significantly preserved in GSK-LSD1-treated mice subjected to permanent coronary ligation (Figure 2A). The same beneficial and significant effect of GSK-LSD1 on ventricular function was observed in hearts with transient coronary ligation (Figure 2B).

**Figure 2.**
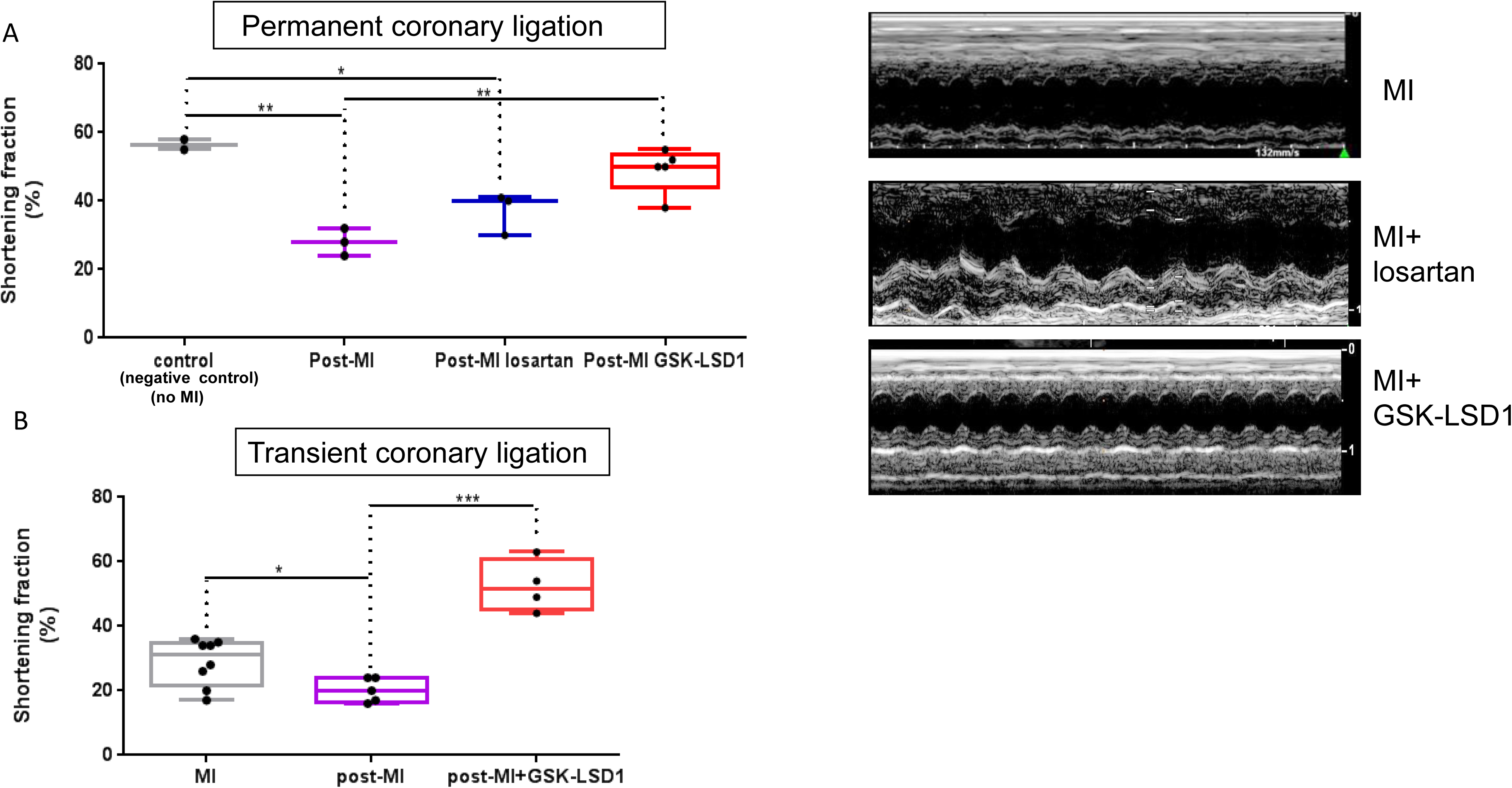
Shortening fraction of mouse hearts under control condition (negative controls are healthy mice with no-MI and no treatment) or after permanent (A) or transient (B) coronary ligation (post-MI). Mice after permanent coronary ligation were treated with losartan (n=4) or GSK-LSD1 (n=5) and mice with transient coronary ligation with GSKLSD1 (n=4). The right panels show some representative echocardiography recordings under the M-mode. Significantly different: *p<0.05, ** p< 0,01,***p<0,001, n=5 mice.

We further tested whether a process of neo-angiogenesis could occur following the clearing of fibrosis. Heart sections were stained with anti-smooth muscle actin (SMA. SMA+ vessels were scored. Figure 3 illustrates that hearts from MI mice treated with GSK-LSD1 featured much many coronary vessels than hearts from non-treated MI mice.

**Figure 3:**
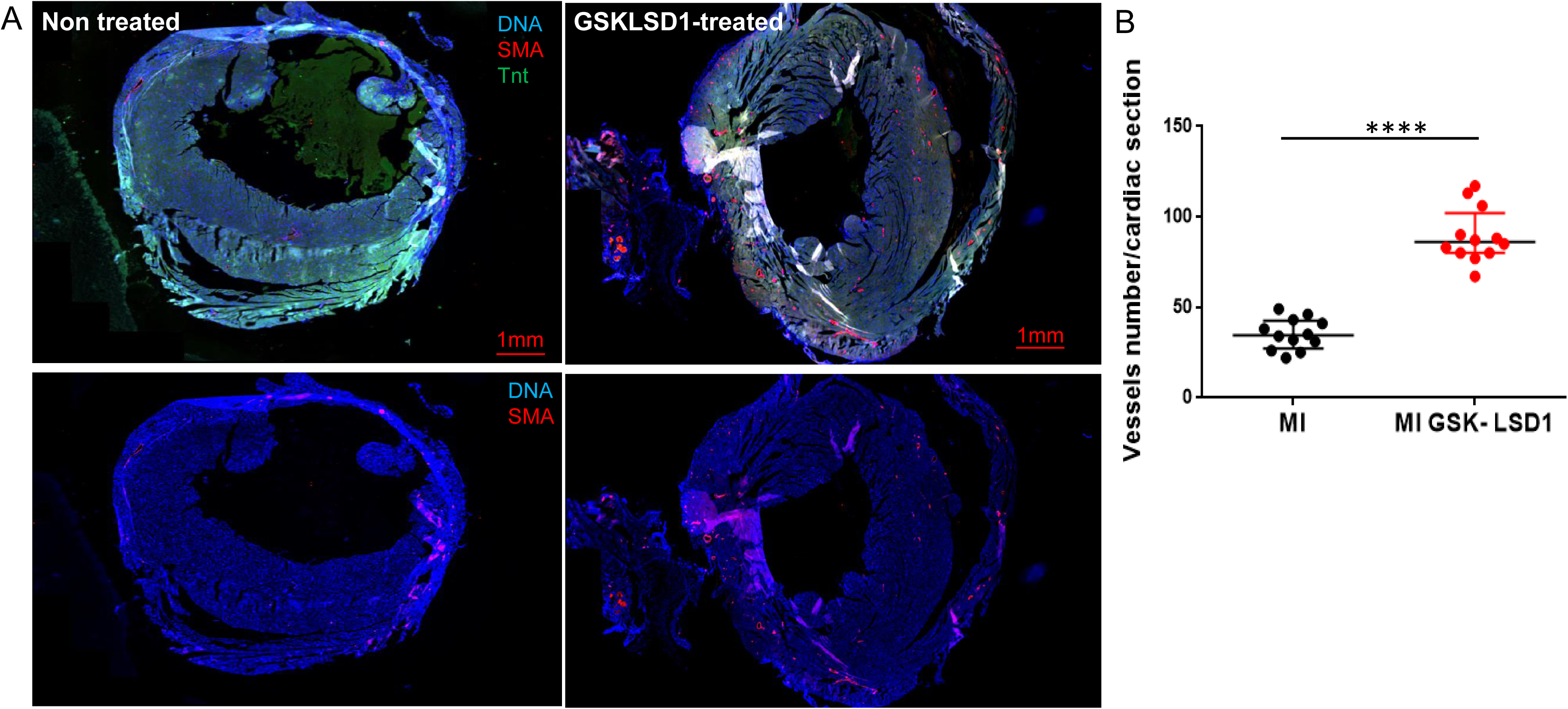
Angiogenesis in post-MI mouse hearts. (A) Cardiac sections of post-MI mouse hearts (treated with LSD1 inhibitor (i) or not (non-treated) were stained with anti SMA, anti-TnT and dapi (DNA). Images were acquired using a scanner (axioscan Zeiss) at 20X magnification and Zen software. (B) SMA+ areas were scored in 3 sections /hearts, n=4 hearts using IMARIS software. Two ways anova test: significantly different p<0.001.

To assess the anti-fibrotic effect of inhibition of LSD1 in a more clinically relevant model, we used a pig model of repaired tetralogy of Fallot which featured right ventricular interstitial fibrosis(18). Pigs were treated with the inhibitor (1.5 mg/Kg) incorporated in the food 3 weeks (median 21 days) after the surgery for a median duration of 30 days. Echocardiography was performed at the time of euthanasia 6 months post-surgery.

Figure 4 illustrates the extent of fibrosis in non-treated and GSK-LSD1-treated pig hearts. As expected, Sirius red staining was extended over the whole ventricle in non-treated pigs. In contrast, collagen staining was more discrete in GSK-LSD1-treated animals (Figure. 4A). The Sirius red staining was confirmed by monitoring the fluorescence emitted by collagen when excited at 560 nm laser in confocal microscopy (Figure 4B). Immunostaining of both periostin, a fibroblast marker and smooth muscle actin ( a marker of myofibroblasts) showed a reduction of both cell types in LSD1 inhibitor treated pig right ventricles ( Figure 4C).

**Figure 4:**
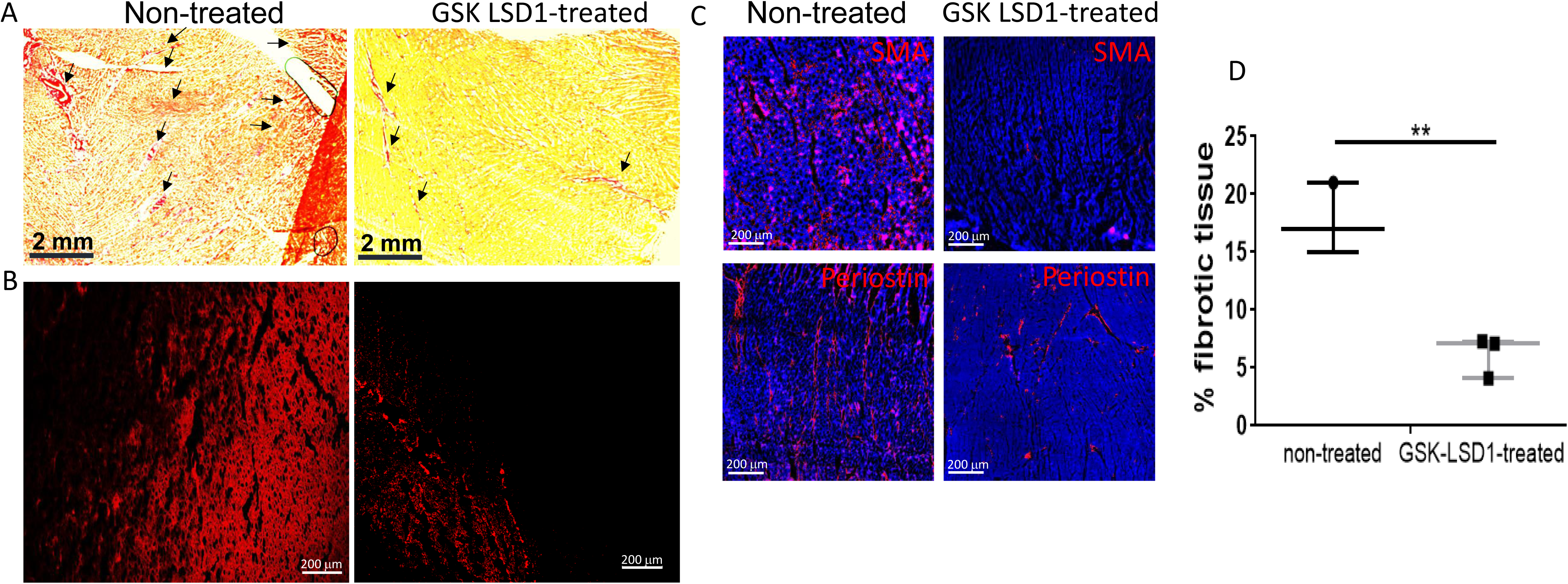
**Fibrosis in rTOF pig right ventricle.** Cardiac sections from the free wall of the right ventricle from non-treated or GSK-LSD1-treated pigs were stained with Sirius red (A) or collagen fluorescence was imaged using confocal microscopy (B). (C) Cardiac sections were also stained with anti-periostin and anti-smooth actin (SMA) to mark fibroblasts and myofibroblasts. (D) The fibrotic area as pointed by arrows in (A) was scored using Image J.

Interestingly in contrast to mice, we did not observe any neo-angiogenesis in pig hearts treated with GSK-LSD1 (data not shown).

Next, we aimed to evaluate the right ventricular function by echocardiography. The RV/LV diameter ratio showed a dilation of the right ventricle that was less pronounced in GSK-LSD1-treated pig hearts than in non-treated pigs hearts. In the same way, normalized end-diastolic RV diameter and area tended to return to normal values in the treated group without reaching significant differences.

The RV wall was significantly less thick in GSK-LSD1-treated heart than in control hearts pointing to les hypertrophy (Figure 5, and supplementary videos).

**Figure 5:**
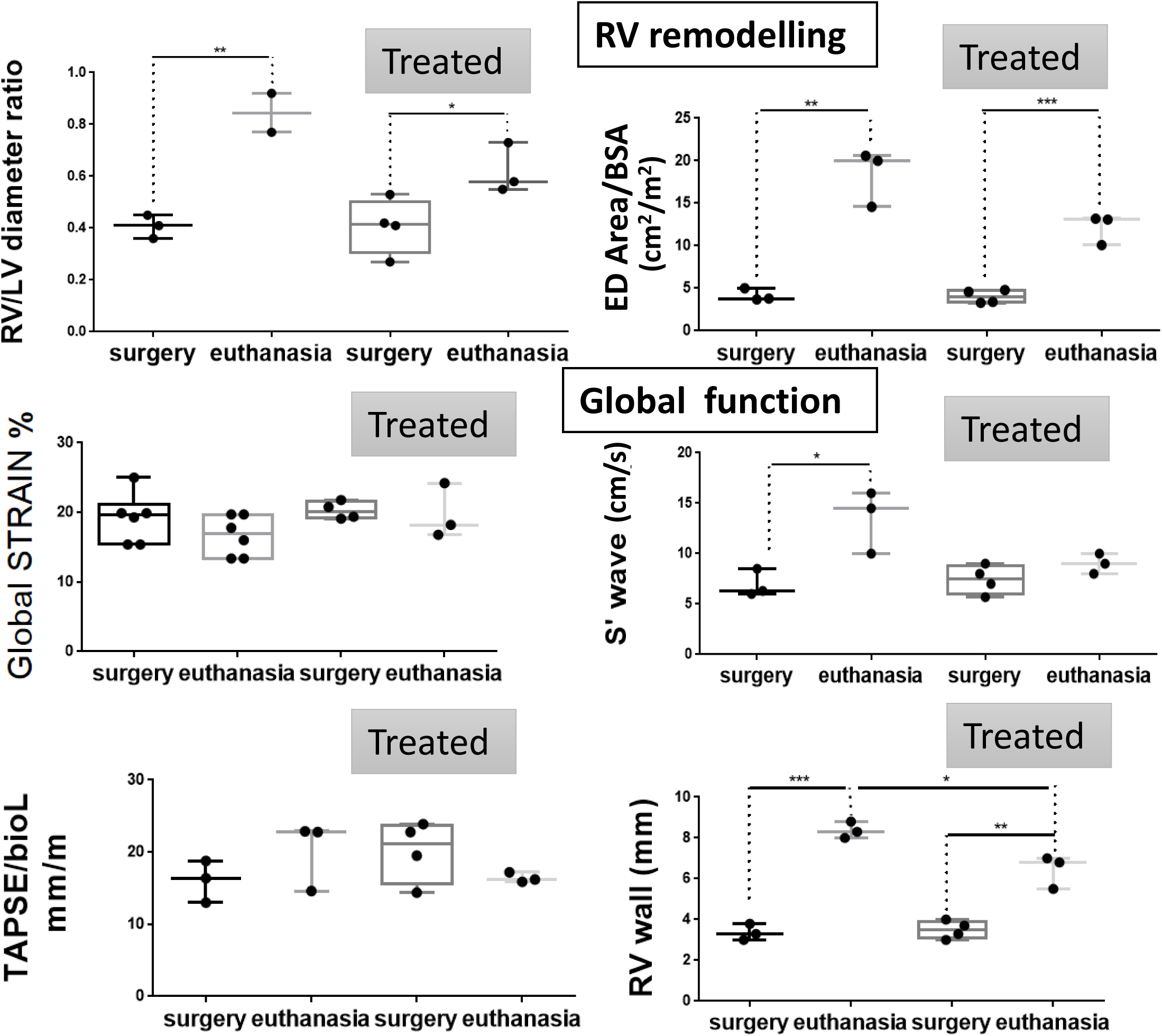
Right ventricular function in pig rTOF. Functional parameters of pigs either non- treated or treated with GSK-LSD1 (Treated) before the surgery (surgery) and at the end point of the experiment (euthanasia). The top graphs reflect the remodeling of the RV and the bottom ones, the function and contractility. The ultrasound images of non-treated (left) or GSK-LSD1- treated pig (right, Treated) show a RV less dilated and with a thinner wall after treatment. T-test was used to compare two time points *p<0.05, ** p< 0,01,***p<0,001.

Changes in normalized TAPSE used to assess global RV dysfunction before, and after development of fibrosis was not significantly and differentially affected in GSK-LSD1-treated pig hearts than in control heart. Nevertheless, on Tissular Doppler imaging, S’ wave tended to return to basal values in the treated group. It should be noted that TAPSE reflects only the longitudinal function of the basal RV free wall. Regarding the RV contractility assessed by strain parameters, no difference of the peak global longitudinal strain was observed between the two groups (Figure 5).

## DISCUSSION

We found that the treatment with LSD1 inhibitors of both mice following myocardial infarction and pig with right ventricular failure significantly reduced fibrosis. Accordingly, the LSD1 inhibitors significantly improved the recovery of cardiac function after MI in mice. The effect of inhibition of LSD1 in the pig model of repaired tetralogy of Fallot was also significant to prevent the formation of fibrotic interstitial tissue. This resulted in limiting both dilation of the ventricle and hypertrophy of the RV. Although in MI mice treated with LSD1 inhibitor, a neo-angiogenesis was observed, this effect cannot be directly attributed to LSD1 inhibition. Indeed this was not observed in pigs also treated with LSD1 inhibitor. Furthermore, LSD1 itself has been reported to be a HIF1 -dependent pro-angiogenic agent (19). This neo-angiogenesis may rather occur more efficiently in rodents as a secondary process following the clearing of fibrosis.

The treatment of pigs in this pilot study was short (3 weeks) which could have limited its impact. However, LSD1 turns out to be a target to significantly limit the deleterious effect of the right ventricle remodeling and in turn to preserve RV function.

LSD1 inhibitors likely target several cell types within the heart including fibroblasts, activated fibroblasts and macrophages. Indeed, besides cardiomyocytes, it has also been reported that a specific downregulation of LSD1 in Angiotensin II-induced myofibroblasts prevented the intracellular upregulation of transforming growth factor β1 as well as, its downstream effectors Smad2/3, p38, ERK1/2 and JUNK(16). LSD1 inhibition also mediates the anti-fibrotic action of Salsolinol(20). While required during embryogenesis, Low LSD1 activity in adult is increased in cardiac diseases(21), which makes the enzyme an appropriate therapeutic target.

Interestingly LSD1 has also been involved in inflammatory processes through immune cells including macrophages(22–24). The inflammation is at the origin of fibrosis and the macrophages at the crossroad of such a process(25). LSD1 inhibitors by targeting several cell types are thus potent modulators of early (i.e. inflammation), middle stage (the EMT process) and late stage (i.e. activation of myofibroblasts) fibrosis.

The molecular mechanism of action of LSD1 inhibitors may be several. LSD1 is together with the REST corepressors (RCORs) the core component of the LSD1/CoREST/HDACs transcriptional repressor complex. It acts at the level of chromatin as a regulator of the transcriptional machinery. The chromatin modifiers complex represses the epithelial or endothelial genes(15) while activating the EMT genes. This explains the pathological effect of LSD1 when reactivated during a pathological situation such as inflammation, cancer or fibrosis. This however does not exclude another potential effect of LSD1 inhibitor independent from the demethylase activity(26).

LSD1 inhibitors are tested in many clinical trials in oncology(27).These are potent blockers of metastasis which share the process of EMT with fibrosis. EMT is also an active process in the epicardium in diseased atria leading to atrial fibrillation(28). LSD1 inhibitors could also be considered to prevent the disease. LSD1 inhibitors may turn out to be potent drugs to treat fibrosis in many organs such as kidney an lungs(29–31), two major clinical burden.

Our study together with others(5) reveal that epigenetic modifiers, and more specifically LSD1 inhibitors could be potent drugs for cardiovascular diseases.

## Limitations of the study

While the pig model is a faithful model in a clinical situation, it features some limitations; The pigs were treated for only 3 weeks and we could not monitor heart function longer than 2 months later. This is due to the fast growing of these animals which prevents their manipulation when they get over 90 kg, what is their weight 8 months after their surgery to engineer the model.

## SUPPLEMENTARY MATERIALS

Figure S1: Individual mouse weight (g) in the course of experiments. Figure S2: LSD1 activity in mouse adult heart. Figure S3: extent of fibrosis in left ventricle of post-myocardial infarction (MI) in mice non treated (red square) and mice treated with the reversible LSD1 inhibitor SP-2509. videos of RV echocardiography from a non-treated- and a LSD1i-treated pig

## AUTHOR CONTRIBUTIONS

DA carried out the experiments, VL coordinated the pig experiments and performed echocardiography, DB helped in pig experiments, AM performed animal surgeries, JG funded part of the work and reviewed the manuscript, MP designed the study and wrote the manuscript.

## INSTITUTIONAL REVIEW BOARD STATEMENT

All animal experiments were carried out according to the European Community guiding principles in animal experimentation. The study was reviewed by ethical committees of the Aix-Marseille and Bordeaux Universities and agreements were obtained from the French Ministère de l‘Agriculture.

