## supplementary figures for "LSD1 demethylase inhibition prevents cardiac fibrosis in both ischemic and congenital diseases in mice and pig models"

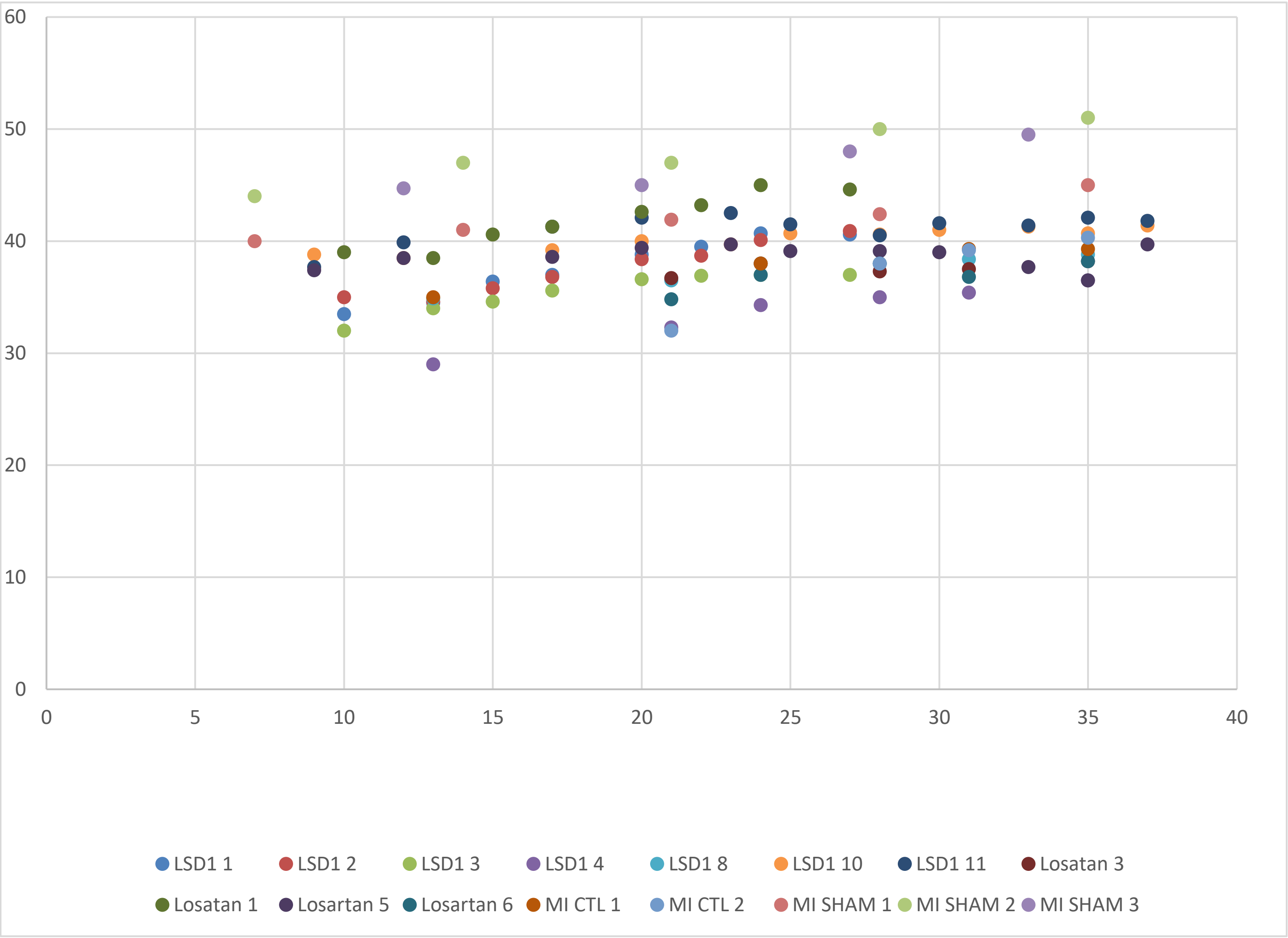


Figure S1:Individual mouse weight (g) in the course of experiments

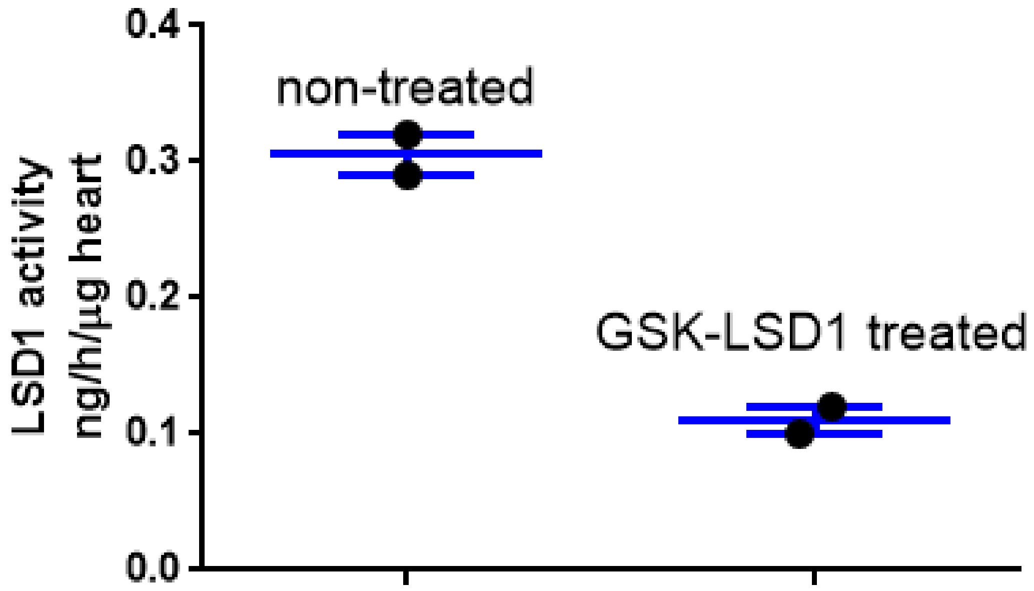


Figure S2: LSD1 activity in mouse adult heart

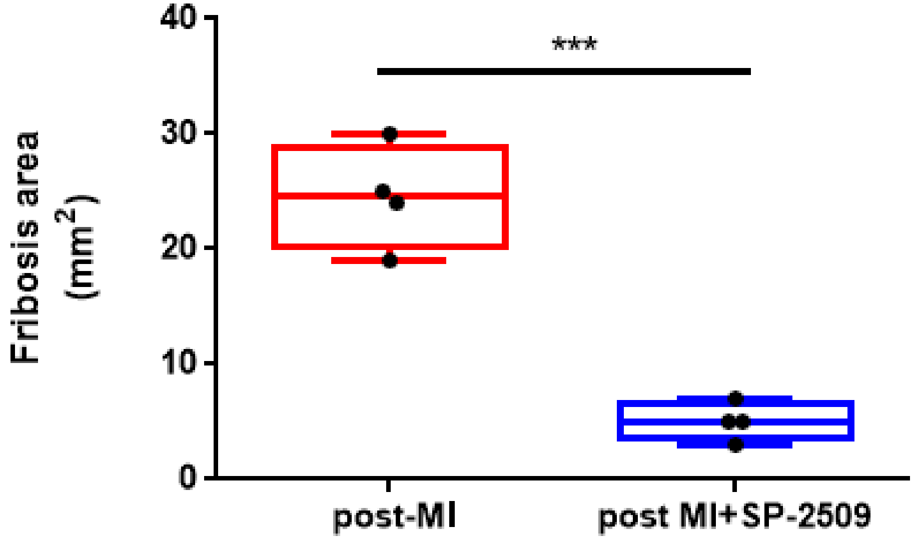


Figure S3: extent of fibrosis in left ventricle of post-myocardial infaction (MI)
In mice non treated ( red square) and mice treated with the reversible LSD1 inhibitor SP-2509
